# Transcriptomic Resilience of a Coral Holobiont to Low pH

**DOI:** 10.1101/157008

**Authors:** Raúl A. González-Pech, Sergio Vargas, Warren R. Francis, Gert Wörheide

**Author notes:** Present address: Institute for Molecular Bioscience, The University of Queensland, 9Brisbane, QLD 4072, Australia. Correspondence: Gert Wörheide.

## Abstract

Ocean acidification is considered as one of the major threats for coral reefs at a global scale. Marine calcifying organisms, including stony corals, are expected to be the most affected by the predicted decrease of the surface water pH at the end of the century. The severity of the impacts on coral reefs remains as matter of controversy. Although previous studies have explored the physiological response of stony corals to changes in pH, the response of the holobiont (i.e. the coral itself plus its symbionts) remains largely unexplored. In the present study, we assessed the changes in overall gene expression of the coral *Montipora digitata* and its microalgal symbionts after a short (three days) and a longer (42 days) exposure to low pH (7.6). The short-term exposure to low pH caused small differences in the expression level of the host, impacting mostly genes associated with stress response in other scleractinians. Resilience to Acidification of a Coral Holobiont Longer exposure to low pH resulted in no significant changes in gene expression of the coral host. Gene expression in the eukaryotic symbionts remained unaltered at both exposure times. Our findings suggest resilience, in terms of gene expression, of the *Montipora digitata* holobiont to pH decrease, as well as capability to acclimatize to extended periods of exposure to low pH.

## Introduction

Ocean acidification, the decreasing pH of ocean surface waters, is considered a global threat to coral reefs (Hoegh-Guldberg et al., 2007). A Business-as-Usual (BaU) carbon emissions scenario predicts surface water pH should reach values ranging between 7.8 and 8.0 by the end of the century, likely posing a challenging future for marine calcifying organisms (Kleypas et al., 2005; Jokiel et al., 2008; De'ath et al., 2009). Among these organisms, scleractinian corals have received particular attention due to their pivotal role as reef builders (Shinzato et al., 2011). In this regard, ocean acidification has the potential to decrease reef-accretion rates and negatively impact these important ecosystems that serve as biodiversity reservoirs, allow the development of other marine ecosystems, and provide important economic and environmental services (Moberg and Folke, 1999; Chen et al., 2015).

The mechanisms by which exposure to acidified water affect calcium carbonate accretion in corals appears to be related to the maintenance of pH-homeostasis at the tissue-skeleton interface, which allegedly makes calcification more energetically expensive (Krief et al., 2010; McCulloch et al., 2012; Venn et al., 2013; Holcomb et al., 2014; Cyronak et al., 2015, Von Euw et al., 2017). Other observed physiological changes triggered by low pH include impaired reproduction, growth and metabolic functions, and anomalies in skeletal morphology (Albright et al., 2008; Albright et al., 2010; Morita et al., 2010; Suwa et al., 2010; Albright and Langdon, 2011; Nakamura et al., 2011). Symbiosis can also be affected as coral bleaching can be triggered by ocean acidification (Anthony et al., 2008). In addition, decreased photosynthetic productivity has been reported in *Symbiodinium* (the microalgal symbiont inhabiting corals) exposed to low pH water, and a delayed establishment of symbiosis has been observed in coral larvae exposed to this condition (Anthony et al., 2008; Crawley et al. 2010). However, the impacts of exposing corals to low pH vary across species and life stages (Kaniewska et al., 2012; Moya et al., 2012; Moya et al., 2015; Davies et al., 2016).

Previous findings suggest that corals can tolerate low pH by using different strategies, for example, through the reduction of their metabolic rate and oxygen consumption (in larvae) or by increasing calcification rates (in adults) (Nakamura et al., 2011; Rodolfo-Metalpa et al., 2011). Better characterization of the responses of these organisms to ocean acidification is therefore fundamental in order to better understand the potential of scleractinian corals to cope with the challenges associated with a changing environment. Within this framework, we sequenced the transcriptome of *Montipora digitata,* a coral commonly found in seawater aquaria and available in culture in different countries, and of its symbiotic microalgae and assessed the changes in global gene expression as result of exposing *M. digitata* to low (7.6) pH water for 3 and 42 days. To date, most studies have focused on the physiological response of stony corals to changes in pH, leaving the coupled transcriptional response of corals and their symbionts to ocean acidification largely unexplored. Here we show that the *M. digitata* holobiont is remarkably tolerant to low pH water exposure, highlighting the potential for resilience and acclimation of corals to ocean acidification.

## 2 Methods

### 2.1 Experimental setup and biological material

The experimental setup consisted of two tanks (control and treatment), each of them containing 20 L of artificial seawater coming from one of the (360 L) aquarium systems available at the Molecular Geo‐ & Palaeobiology Laboratory of the Department of Earth and Environmental Sciences, Palaeontology & Geobiology, LMU München (Supplementary Fig. S1). Water in the experimental tanks was replaced 3 to 5 times per hour with water from the main (360 L) tank. To simulate ocean acidification conditions, a pH-electrode (LE4099, Mettler Toledo) connected to a pH computer was used to control the injection of CO_2_ into the water of the treatment tank to be kept pH at 7.6 (treatment tank). In each experimental tank a water pump set to a flow rate of 300 L/h was used to keep the water circulating and to better mix the CO_2_ injected into the treatment tank. Water returning from the treatment tank was equilibrated to pH ∼8.0 by letting it flow through a 25 cm column containing limestone grains with diameters ranging from 2 to 5 mm and simultaneously injecting air (300 to 500 L/h). After this, the treated water was mixed with the remaining water in the main (360 L) tank. Both tanks were illuminated using a Mitras LX 6200-HV lamp simulating tropical light cycles; illuminance ranged between 19 and 22 kilolux at the seawater surface of both tanks. During the experiment, water pH in each experimental tank was controlled every day and was 9 lrecorded every hour from the second day onwards with a pH electrode LE4099 Mettler Toledo and a pH-meter PCE-PHD 1. During the experiment, the pH of the treatment tank was relatively constant (mean = 7.64, sd = 0.02), while the pH in the control aquarium fluctuated between 7.90 and 8.26 according to the respiration/photosynthesis cycle of corals and symbionts (mean = 8.08, sd = 0.11).

Coral nubbins (n = 20) were obtained from branches of two adult *Montipora digitata* colonies from our coral culture stock. The individual explants were cultured for over 2 months in the main (360 L) aquarium under the same light conditions used for the experiment.

Approximately one month before starting the pH experiment, the nubbins were transferred to the control aquarium of the experimental setup (see above) for acclimation. Afterwards, half of the corals were moved to the treatment tank and sampling was conducted after 3 and 42 days of exposure. Five corals were sampled from each tank at each sampling time.

### 2.2 RNA extraction and sequencing

At each sampling point, the nubbins were flash-frozen in liquid nitrogen, broken in smaller pieces with mortar and pestle (always in liquid nitrogen) and stored at -80 °C until further processing. For RNA extraction, samples were homogenized in TRIzol Reagent (Thermo Fisher Scientific) with a SilentCrusher M and a Dispersion Tool 8 F (Heidolph) and total RNA was extracted using a slightly modified version of the Chomczynski method (Chomczynski and Sacchi, 1987). RNA quality and concentration were controlled with a NanoDrop ND-1000 spectrometer and with a BioAnalyzer 2100 (Agilent Technologies, Santa Clara, CA, USA). An additional cleaning step with Agencourt RNAClean XP Magnetic Beads was done if required. RNA concentration and quality requirements for library preparation and sequencing were achieved for all samples obtained after the first sampling time, for the second sampling time only 6 samples (3 control + 3 treatment) met quality requirements and were further used. Library preparation and sequencing (50bp PE; HiSeq2500) was done at the GeneCore of the European Molecular Biology Laboratory (EMBL) in Heidelberg, Germany. The reads generated for the second sampling time were strand-specific and further used for *de novo* transcriptome assembly. The reads generated were uploaded to the Short Read Archive of the European Nucleotide Archive under the project accession PRJEB21531 121(athttp://www.ebi.ac.uk/ena/data/view/PRJEB21531).

### 2.3 Transcriptome assembly and annotation

Illumina read pairs were quality controlled with FastQC v0.63 (Andrews, 2010), low quality bases and reads were removed with Trimmomatic v0.32 (Bolger et al., 2014). The surviving pairs were further processed with the program Filter Illumina v0.40 of the Agalma/Biolite suite (Dunn et al., 2013) to ensure the removal of adapters and low quality reads. Reads from putative bacterial contaminants were filtered out by aligning the surviving pairs against all bacterial genomes on RefSeq as of July 2015 (ftp://ftp.ncbi.nlm.nih.gov/genomes/refseq/bacteria) and discarding mapping pairs.

De novo transcriptome assembly was done with Trinity v2.0.6 using only strand-specific sequences *(i.e.* the surviving read pairs from the second sampling time) (Grabherr et al., 2011). The assembled contigs from coral and dinoflagellates were separated with Psytrans (https://github.com/sylvainforet/psytrans) using the *Acropora digitifera* v0.9 (Shinzato et al., 2011) and *Symbiodinium minutum* Clade B1 v1.0 (Shoguchi et al., 2013) predicted proteins as references. The minicircle sequences of *Symbiodinium* sp. Chloroplast (Barbrook et al., 2014) were recovered manually from the contigs assigned to the coral. Coding sequences (CDS) from the assembled transcripts were predicted with TransDecoder v2.0.1 (https://transdecoder.github.io). Only transcripts with predicted proteins were used as the reference transcriptome for both coral and symbionts. Completeness of the transcriptomes was assessed by search of the sequences against the Core Eukaryotic Genes Dataset (CEGMA) (Parra et al., 2007; Francis et al., 2013). Data processing and transcriptome assembly were mostly executed on the Molecular Geo‐ and Paleobiology Lab’s Galaxy server (Goecks et al., 2010). Differentially expressed genes (DEGs hereafter) were annotated by BLASTN *(E* = 10^-10^) against the Non-Redundant (nr) database of the NCBI and by BLASTX (*E* = 10^-10^) against the SwissProt database (both downloaded in February 2015). The best SwissProt hit of each DEG was used to retrieve its associated Gene Ontology terms through the QuickGO online tool (https://www.ebi.ac.uk/QuickGO). In addition, predicted proteins of the DEGs were annotated with the protein identifiers of the *Acropora digitifera* v0.9 and *Symbiodinium minutum* Clade B1 v1.2 by BLASTP (*E* = 10^-10^). A Galaxy history containing some of the steps previously mentioned can be accessed at (XXXXXXX). The transcriptome annotations can be found in https://github.com/PalMuc/Montipora_digitata_resources.

### 2.4 Differential gene expression analysis

For each sampling time, the remaining reads from each sample were mapped with Bowtie2 (Langmead et al., 2009) to the reference transcriptomes of *M. digitata* and *Symbiodinium* sp. Read counts from isoforms groups were added to derive a matrix of counts per Trinity-component. Differences between conditions and colonies were assessed with the *adonis* function of the *vegan* R library (Oksanen et al., 2013). As in other ocean acidification studies (Davies et al., 2016; Kurman et al., 2017), the sample-dependence effect caused by one single tank per condition could not be corrected (Riebesell et al., 2011; Cornwall and Hurd, 2016; but see Oksanen, 2001 and Schank and Koehnle, 2009 for a discussion on pseudoreplication). DEGs between conditions were detected using the package DESeq2 v1.8.1 (Love et al., 2014). DEGs were selected by their significant change in expression as assessed by a Wald test; P-values were adjusted with the Benjamini-Hochberg correction (Benjamini and Hochberg, 1995). Gene Ontology (GO) terms enrichment was assessed with topGO v1.0 (Alexa and Rahnenfuhrer, 2010) in R v3.2.1. Protein domains of the DEG products were identified using PfamScan and PFAM A release 30.0 (both downloaded from ftp://ftp.ebi.ac.uk/pub/databases/Pfam/). The counts and the R scripts used to analyze the data can be found in the project repository (https://github.com/PalMuc/Montipora_digitata_resources).

## 3 Results

### 3.1 Reference transcriptomes

The *Montipora digitata* meta-transcriptome was assembled using over 190 million of strand-specific read pairs. The transcriptome assembly yielded 179,298 sequences, of which 123,710 (69 %) were assigned to the coral host and 55,588 (31 %) to the symbiont. These transcript sets resulted in 41,852 and 34,057 predicted CDS for the coral and symbionts, respectively. The N50 length was 1,101 bp for the coral and 744 bp for *Symbiodinium* sp. G+C content of both reference transcriptomes were 43.9% and 54.5%, consistent with previous reports for the Order Scleractinia and *Symbiodinium* spp., respectively (Bayer et al., 2012; Shoguchi et al., 2013; Sun et al., 2013; Shinzato et al., 2014). Summary quality statistics for both transcriptomes are summarized in Table 1. Most of the Core Eukarytic Genes (CEGs) were found in the transcriptomes, 89.5% in *M. digitata* and 79.4% in *Symbiodinium* sp. (Supplementary Table S1). In addition, the level of fragmentation of the assembled transcriptomes appears to be low as judged by the low number of CEGs that are longer than the queries.

**Table 1.** Summary quality statistics for the reference transcriptomes of the M. digitata and Symbiodinium sp.

| Statistic | *Montipora digitata* Transcriptome | *Symbiodinium* sp. Transcriptome |
| --- | --- | --- |
| GC content | 43.9 % | 54.5 % |
| N50 (bp) | 1,101 | 744 |
| Max. length (bp) | 10,503 | 6,318 |
| Mean length (bp) | 872 | 648 |
| Median length (bp) | 645 | 540 |
| Min. length (bp) | 297 | 297 |
| Total length (bp) | 36,510,222 | 22,095,753 |
| Num. of sequences | 41,852 | 34,057 |

Only 5,541 (13.2%) out of the 41,852 genes contained in the transcriptome of *M. digitata* transcriptome could not be annotated with any of the databases. A large portion (48.6%) of transcripts had hits against sequences from all of the three databases used and about 15% of the total transcripts hit exclusively sequences of the *A. digitifera* genome (Fig. 1). On the other hand, most of the sequences in the transcriptome of *Symbiodinium* sp. were annotated with genes from the *S. minutum* genome (85.5% of all transcripts), and almost half of them 195(48.4%) had hits exclusively against this genome. Similarly to the coral transcriptome, only a small fraction of the transcripts (12.9%) lacked any annotation (Fig. 1).

**Figure 1.**
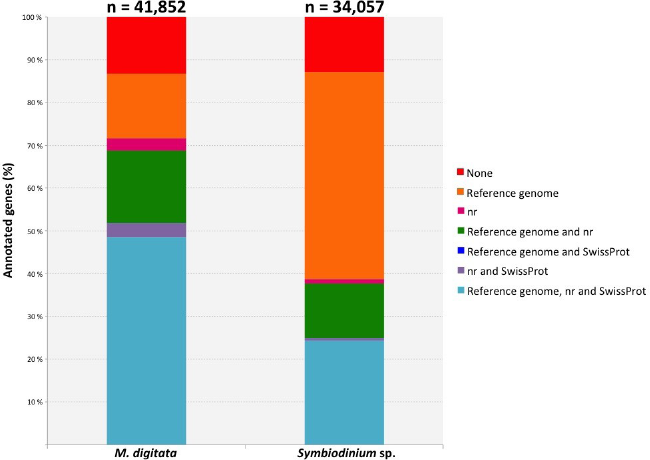
Bar plot displaying different fractions of the annotations for both, coral and symbiont, transcriptomes. The total number of genes for each transcriptome is shown above the corresponding bar. Each color represents an annotation category specified in the legend on the right and the lines connecting the bars indicate the position of the same category for each dataset. The category with the largest fraction in the coral transcriptome corresponds to those genes that had records from the three annotation resources. Genes with no annotation records are shown in red and represent a minor fraction of each transcriptome. Reference genomes used were *Acropora digitifera* (scleractinian coral) and *Symbiodinium minutum* (dinoflagellate symbiont).

### 3.2 Coral response to low pH

No significant differences in global expression were found with the *adonis* exploratory test between conditions and between colonies for neither the short (p_condition_ = 0.40, p_colony_ = 0.19) nor the longer (p_condition_ = 0.73, p_colony_ = 0.76) exposures of the coral to low pH. These findings are also supported by the variance in expression of the coral genes at both sampling times (Fig. 2) and by the variance-corrected counts of mapped reads per gene per sample (Supplementary Fig. S2).

**Fig 2.**
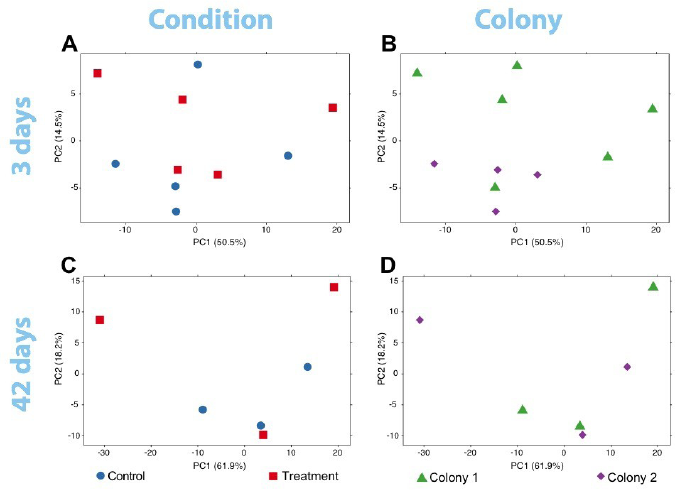
PCA plots of the variance of expressed genes in the coral nubbins (A) between colonies and (B) conditions after three days of exposure to low pH. Variance of gene expression is also plotted after the 42-days exposure comparing conditions (C) and colonies (D). No grouping patterns in any comparison can be distinguished. Dot color and shape codes are displayed at the bottom.

Still, the DeSeq2 gene expression analysis yielded 18 genes with significant (p_adj_ < 0.05) differences in expression level due to exposure to low pH after three days (model design: ∼ colony + condition). The log_2_-fold changes for these genes ranged between -0.70 and 0.73 (Supplementary Table 1). Six up‐ and 12 downregulated genes were found. From the six upregulated genes, two (including a metalloprotease) contain zinc-binding domains and others have RNA/DNA binding domains. Among the downregulated genes, proteins similar to Noelin, to an αB-crystallin and to a “platelet-derived” growth factor were found. A protein with no annotation but containing a domain of the tumor necrosis factor receptor superfamily (TNFR) was also downregulated. GO terms related to organ development, cell differentiation and cell chemotaxis were enriched in this set of genes (Supplementary Table S2). Upregulated genes were rich in GO terms related to the negative regulation of calcium ion-dependent exocytosis, regulation of extracellular matrix assembly, collagen catalysis, changes in bone morphogenesis, negative regulation of cell-substrate adhesion, transmembrane transport of zinc and response to antigenic stimulus. Additionally, the Notch-signaling pathway appeared to be positively regulated, contrary to previous findings (Kaniewska et al., 2012). No DEGs were found in the coral after 42 days of exposure to low pH.

### 3.3 Symbiont response to low pH

The exploratory analyses of gene expression of *Symbiodinium* sp. suggest no change at any sampling time (Fig. 4). The *adonis* permutation test resulted in no significant differences between conditions and colony after the first (p_condition_ = 0.62, p_colony_ = 0.78) or the second 228(p_condition_ = 0.88, p_colony_ = 0.99) exposure period. No DEGs were found at any time.

## 4 Discussion

### 4.1 Mild stress response of the coral to low pH

The present study used RNA-Seq to assess the coupled response of an adult stony coral (*Montipora digitata*) and its microalgal symbionts to low pH. Compared to previous investigations (Kaniewska et al., 2012; Moya et al., 2012; Moya et al., 2015), our results suggest a mild gene expression response of the adult coral holobiont to a three-days exposure to acidified water in terms of number of DEGs and log_2_-fold change. Still, the log_2_-fold changes we found in this study are comparable to those observed in the resilient coral *Siderastrea siderea* under laboratory-induced ocean acidification conditions (Davies et al., 2016). This suggests resilience of *M. digitata* to low pH.

The few DEGs found after short-term exposure of *M. digitata* to low pH (7.6) have been shown to be involved in stress response in corals. Zinc-metalloproteases, for instance, seem to be relevant for heat tolerance in these organisms and might play a role in regulation of apoptosis and cell repair (Barshis et al., 2013). Other stress related genes were found among the significantly downregulated genes, like the αB-crystallin-like protein, an HSP20 family member. αB-crystallin seems to play a role in stress response in other corals (e.g. *Orbicella annularis* and *O. faveolata),* but its exact function is yet unknown (Downs et al., 2002). A TNFR domain-containing protein is another example, and changes in expression levels of proteins carrying this type of domain display have been found in other corals under stress conditions (Barshis et al., 2013; Seneca and Palumbi, 2015; Yuan et al., 2017). Another down‐ regulated gene codes for a protein containing a platelet-derived growth factor domain, though the regulation of this kind of protein upon exposure to low pH has not been previously reported in corals. However, it is already known that homologs of human bone-morphogenesis proteins (BMP2/4) participate in skeleton production of marine calcifiers, including corals (Green et al., 2013). In fact, proteins carrying this domain are currently used for bone regeneration in humans, and its use, in combination with coral skeleton, has been proposed to treat bone defects (Parizi et al., 2012).

According to previous investigations, coral larvae are able to acclimate to acidified water within a few-days time span (Moya et al., 2015). Such a rapid acclimation by the adult coral holobiont would thus explain the lack of change in gene expression after 42 days of acidified water exposure in our experiments, a time by which acclimation would likely have been already completed. Our findings in the long-term response are similar to a previous investigation on response of another adult scleractinian *(Acropora millepora)* to a 37-days exposure to low pH (elevated pCO_2_) that reported absence of change in the expression of target genes associated with calcification and metabolism (Rocker et al., 2015). In agreement with this, previous studies suggest that temperature stress has a greater impact in gene expression of corals than pH (Mayfield et al., 2014; Davies et al., 2016), although resilience to stress conditions varies from one species to another (Loya et al., 2001).

### 4.2 Symbiont response

Physiological response to stress conditions does not only depend on coral host resilience but also on the interaction with its symbionts (Hoadley et al., 2015). In the present study, *Symbiodinium* sp. did not seem to be affected by the decrease of pH, possibly because of the more stable environment provided by the coral tissue, which acts as some kind of protection for the algae (Banaszak and Trench, 1996). However, differences in response between *Symbiodinium* spp. *in hospite* and isolated from the host to environmental stressors remain largely unexplored.

## 5 Conclusions

Here we provide baseline data in the form of a reference transcriptome of the scleractinian coral *Montipora digitata* and its dinoflagellate symbiont. We used these data to assess changes in overall gene expression of the different components of the coral holobiont, i.e., the coral host and its microalgal symbionts after different periods of exposure to low pH (7.6) seawater. While the coral host showed an initial stress response after short-term exposure to low pH, an acclimatization of gene expression levels was indicated after longer exposure (42 days) to low pH, whereas its symbionts remained unaffected independent of exposure times. While we cannot exclude tank effects due to the experimental set-up, these findings nonetheless suggest a potential of the *Montipora digitata* holobiont for acclimatization and resilience to lowered seawater pH, and that temperature may play a greater role in coral stress than pH. Both the reference transcriptome we provide and the working hypothesis of acclimatization and resilience in this model system serve as a valuable baseline for future in-depth experiments.

## 5 Author Contributions

GW and SV conceived the study, SV coordinated the lab and analytical work, RGP carried out the experiments, lab work and analysed the data, WRF contributed to bioinformatic analyses, GW acquired the funding.

## 7 Acknowledgments

This study would not have been possible without the support of Gabi Büttner, Simone Schatzle and of Dr. Peter Naumann. Dr. Michael Eitel provided important discussions about data analysis. We would also like to thank Dr. Helmut Blum, Dr. Stefan Krebs and Andrea Klanner, LAFUGA, Gene Center LMU München, for the support during early stages of laboratory work. We acknowledge Dr. Sylvain Foret for his help with Psytrans. SV is indebted to N. Villalobos Trigureros, M. Vargas Villalobos, S. Vargas Villalobos and S. Vargas Villalobos for their constant support.

## Supplementary Material

**Fig S1.**
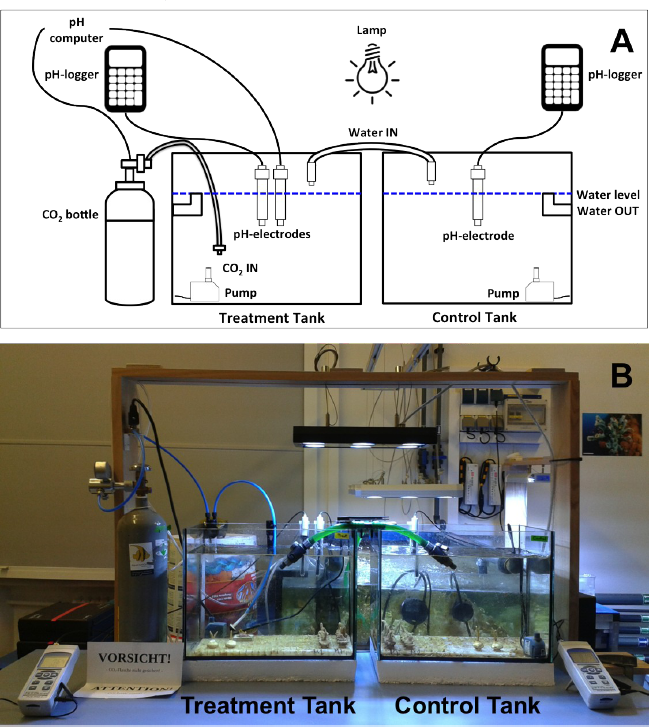
**(A)** Diagram of the aquaria setup. To the left is the treatment tank (pH = 7.6) and to the right the control tank (pH = 8.0 pH). Water from one of the main water systems of the aquarium room flowed into the tanks (water IN) and back again to the big aquarium (water OUT). The acidified water returning to the big aquarium from the treatment tank was treated by letting it flow through a 25 cm column containing limestone grains with diameters ranging from 2 to 5 mm and injecting air with a flow rate from 300 to 500 L/h to cast out the carbon dioxide. pH of the treatment tank was kept at 7.6 by an automated mechanism that injected CO_2_ from a bottle whenever the pH went above this value. During the experiment, pH was recorded with data loggers plugged to pH-electrodes in each tank. Water pumps kept recirculating the water, particularly important for the well mixing of CO_2_ in the treatment tank. A Mitras LX 6200-HV lamp simulating the natural daily light cycles in the tropics was placed above and between the two tanks such that the illuminance was equally distributed. **(B)** A photograph of the actual aquaria setup.

**Fig. S2.**
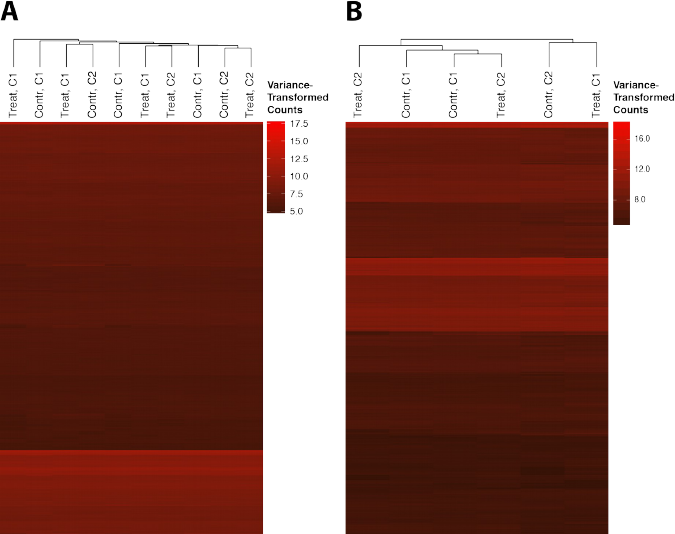
Heatmaps of the transformed counts (using the variance stabilizing normalization of DESeq2) of mapped reads per gene of the coral host at the first **(A)** and second **(B)** sampling times. Each column represents a sample and each row a gene. Dendrograms at the top of the plots cluster the samples by average. Genes are sorted by decreasing variance from top to bottom. Most of the genes with the highest expression values in the coral after three days of exposure to low pH were amongst the least variable **(A)**.

**Table S1.** Completeness of reference transcriptomes based on comparison with the 248 genes in the Core Eukaryotic Genes Dataset (CEGMA). CEG: Core Eukaryotic Genes.

| Category | Coral transcriptome | Symbiont transcriptome |
| --- | --- | --- |
| CEG with hits | 222 | 197 |
| High-confidence full-length matches | 150 | 104 |
| Probable full-length matches | 7 | 5 |
| CEG slightly shorter than query | 5 | 2 |
| CEG much shorter than query | 31 | 33 |
| Probable miss-assemblies | 25 | 51 |
| CEG much longer than query | 4 | 2 |
| CEG with no hits | 26 | 51 |

**Table S2.** DEG in the coral host after a three-days exposure to low (7.6) pH, including gene group identifier, log_2_-fold change, corrected p-value, protein domains found in predicted proteins and corresponding annotation.

| Isogroup | Log<sub>2</sub>-FC | P<sub>adj</sub> | Pfam Domain(s) [E-val] | Annotation |
| --- | --- | --- | --- | --- |
| TR57464\|<br>c0_g1 | 0.73 | 0.00025865<br>7 | Astacin [1.7e-64] | Zinc metalloproteinase nas-14 |
| TR3854\|c8_g1 | -0.70 | 0.00021718<br>1 | OLF [2.2e-33] | Noelin |
| TR12191\|<br>c4_g3 | -0.64 | 0.00378190<br>9 | NA | NA |
| TR55829\|<br>c4_g3 | 0.60 | 0.02163614<br>9 | NA | NA |
| TR8304\|<br>c10_g2 | -0.59 | 0.02044362<br>1 | NA | NA |
| TR25513\|<br>c0_g1 | 0.57 | 0.02286719<br>2 | DUF4772 [9.3e-22],<br>53-BP1_Tudor [4.4e-05] | Zinc finger protein 395 |
| TR5422\|c1_g1 | -0.56 | 0.02163614<br>9 | NA | NA |
| TR47269\|<br>c9_g2 | -0.56 | 0.02044362<br>1 | Rve [5.8e-21] | Uncharacterized protein K02A2.6 |
| TR17622\|<br>c0_g2 | -0.55 | 0.02492151<br>7 | Ets [8.1e-35] | ETS domain-containing protein Elk-1 |
| TR4348\|c4_g1 | -0.54 | 0.04991696<br>9 | NA | NA |
| TR50981\|<br>c7_g3 | -0.53 | 0.02044362<br>1 | TNFR_c6 [1.9e-04] | NA |
| TR60375\|<br>c5_g4 | -0.52 | 0.04926036<br>7 | HSP20 [4.6e-19] | Heat shock protein Hsp-16.1/Hsp-16.11, Alpha-crystallin B chain |
| TR21650\|<br>c1_g1 | -0.50 | 0.01680226<br>2 | NA | Protein with a platelet-derived and vascular endothelial growth factors (PDGF, VEGF) family domain |
| TR28829\|<br>c0_g1 | -0.46 | 0.04926036<br>7 | ELFV_dehydrog_N [3.4e-51] | NADP-specific glutamate dehydrogenase |
| TR25128\|<br>c7_g1 | 0.46 | 0.01680226<br>2 | NA | NA |
| TR1763\|c4_g1 | -0.42 | 0.02286719<br>2 | RRM_1 [2e-10] | Peroxisome proliferator-activated receptor gamma coactivator-related-like protein |
| TR1853\|c1_g2 | 0.40 | 0.02044362<br>1 | NA | Protein with RNA/DNA binding domains |
| TR21661\|<br>c1_g1 | 0.38 | 0.03404108<br>1 | Helicase_C [1.1e-47] | Werner syndrome ATP-dependent helicase-like protein |

